# Heterozygote calling in significantly inbred populations

**DOI:** 10.1101/481101

**Authors:** Rohan Shah, Alex Whan

## 1 Introduction

Realising the potential of large genetic resources requires the ability to perform genotyping efficiently. In populations with tens or hundreds of thousands of SNP markers, it is infeasible to assign genotypes manually; this must be done automatically. This problem is generally solved by the application of mixture models [Xiao et al., 2007, Teo et al., 2007].

We describe a marker calling method able to correctly identify marker heterozygotes, in large populations generated according to some experimental design; e.g. the Collaborative Cross [Threadgill and Churchill, 2012] or MAGIC [Huang et al., 2015]. Identifying marker heterozygotes is of substantial value in these populations, which are substantially inbred with a small amount of residual heterozygosity. If genetic lines can be identified which are heterozygous at a QTL locus, these lines can be used to develop a follow-up population suitable for fine-mapping of that QTL. However, as the amount of residual heterozygosity is low, these lines cannot be identified by an unsupervised method.

We describe a model-based clustering method for identifying marker heterozygotes in significantly inbred populations. Like Xiao et al. [2007] and Teo et al. [2007], our method is also based on mixture models. Unlike those methods, we include a constraint that the heterozygote cluster lies between the two marker homozygote clusters (Figure 1). This constraint accurately represents the structure of the heterozygote cluster in the case of data generated by the Illuimina Infinium genotyping platform. The proposed model-based clustering algorithm was motivated by an eight-parent MAGIC population in spring bread wheat, genotyped using the Illumina Infinium genotyping platform, and we demonstrate its use in that population. The proposed method is implemented in the R package magicCalling.

**Figure 1:**
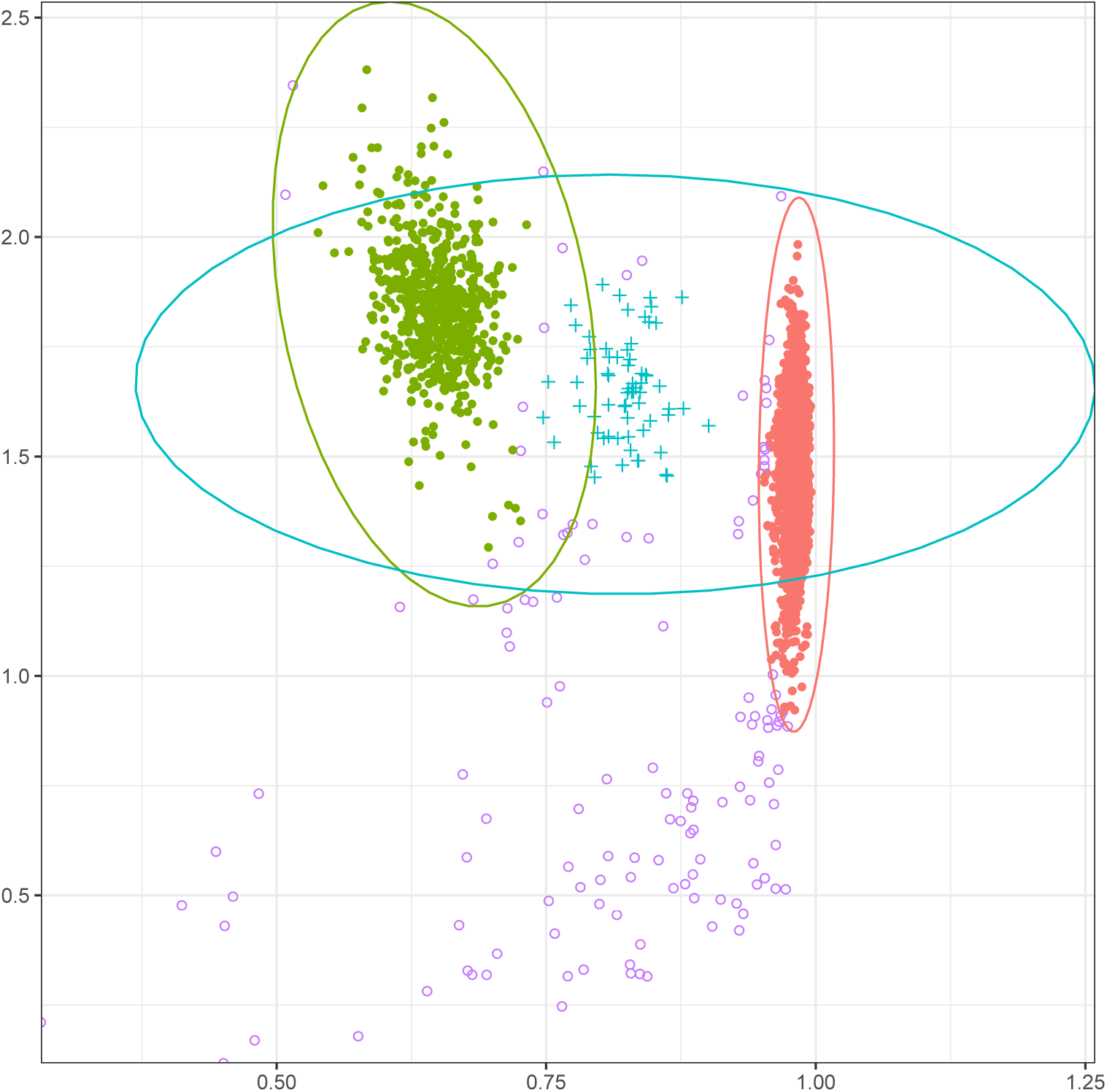
Example of the hierarchical Bayesian clustering approach, applied to data from an 8-parent MAGIC population. Filled points represent lines classified as homozygotes. Crosses represent lines classified as heterozygotes. Hollow points represent outliers, or points not otherwise classified. Ellipses represent covariance matrices of components.

## 2 Model

Assume that we have genotyping data at a single marker, for *N* individuals. This data takes the form of normalized hybridization intensities, which can be represented in a number of different coordinate systems. We assume every hybridization intensity is represented as a strength *r* and a contrast *θ*. As an example, every point in Figure 1 is a hybridization intensity.

Our model of this bivariate data is a four-component normal mixture model, with a hierarchical prior. The first two components correspond to the homozygous marker alleles, and have bivariate normal distributions N (***µ***_1_, **∑**_1_) and N (***µ***_2_, **∑**_2_). The third component represents the marker heterozygotes, and has distribution 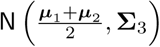; that is, the third component lies between the two heterozygote clusters, and does not have its own independent mean component, although it does have its own covariance matrix. The fourth component represents errors, and has distribution N (***µ***_4_, **∑**_4_). It is intended to be much flatter, and capture all points not represented in the first three components.

We now specify the priors for the mean parameters ***µ***_1_, ***µ***_2_, ***µ***_4_ and covariance matrices **∑**_1_, …, **∑**_4_ using multivariate normal and Wishart distributions. The priors for the mean parameters are

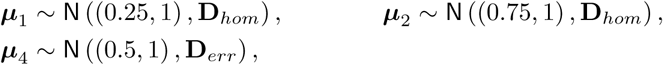

where ***µ***_1_, ***µ***_2_ and ***µ***_4_ are independent. Matrix **D**_*hom*_ is a hyperparameter for the homozygote clusters, and **D**_*err*_ is a hyperparameter for the error cluster. The priors for the covariance matrices **∑**_1_, …, **∑**_4_ are Wishart distributions:

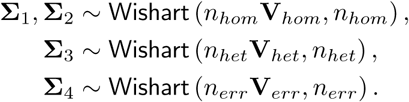

The matrix parameters in these distributions are the expected values of the covariance matrices. So for example, 𝔼 [**∑**_1_] = **V**_*hom*_, and **V**_*hom*_ represents the expected shape of the covariance matrix. Similarly for **∑**_2_, **∑**_3_ and **∑**_4_. The real parameters *n*_*\**_ represent the degree to which the covariance matrix distribution is concentrated around its expectation; a large value indicates a distribution highly concentrated around the matrix parameter. A small value indicates a flatter distribution, and greater a priori uncertainty about the shape of one of the clusters.

The last prior to be specified is the prior on the mixing proportions **p** = (*p*_1_, *p*_2_, *p*_3_, *p*_4_) of the four components. Our prior knowledge is that most lines will be homozygotes, due to inbreeding, and relatively few lines will be heterozygotes or errors. This means we have a prior belief that *p*_1_ + *p*_2_ should be much larger than *p*_3_ + *p*_4_. However *one of p*_1_ or *p*_2_ may be small, as the marker allele distribution is unknown. For example, in our target eight-parent MAGIC population, we will quite often have *p*_1_ ∼ 0.125 and *p*_2_ ∼ 0.875.

To account for this, we define our prior as follows. Let *p*_*hom*_ be the proportion of homozygotes, and let

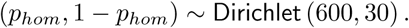

That is, most lines are homozygotes. Let 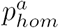 be the proportion of homozygotes that fall into the first cluster, and let

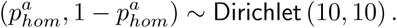

That is, there is no prior belief that one homozygote cluster is bigger than the other. Let *p*_*het*_ be the proportion of heterozygotes, among all lines that are *either heterozygotes or errors*, and let

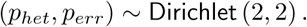

That is, our prior belief is that any line that is not a homozygote is equally likely to be an error or a heterozygote. Combining all these priors, the mixing proportions are

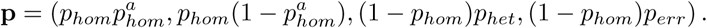

## 3 Implementation

Previous mixture model approaches have used the EM algorithm [Teo et al., 2007]. Our choice of conjugate priors has the advantage that standard software tools can sample from the the posterior distribution. We use the JAGS (Just Another Gibbs Sampler) package [Plummer, 2015] to perform Gibbs sampling for the model described in Section 2. Technically, our use of Gibbs sampling means that the resulting Markov Chain mixes poorly, as the two homozygote clusters will never swap. However, given our clustering context, this type of mixing is irrelevant.

For multi-parent experimental designs, the number of lines is generally large enough that the posterior variance for the model parameters is small, so significantly different parameter estimates from Markov chains with different starting points indicates MCMC non-convergence. This holds in particular for our motivating population, which has 3,412 lines. Because of this, our approach is to run the Gibbs sampler for a certain number of steps, and to use the parameter values at the final step of the Markov chain as estimated parameters.

Given a large enough number of markers, and therefore applications of the model, it is inevitable that some Markov chains will fail to converge. There will also be cases where the model is inappropriate, as the marker is either monomorphic, or has more than two alleles. These cases can be detected using heuristics.

To determine whether a fitted model is acceptable, it is useful to consider:

1. 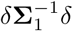 and 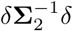, where *δ* = ***µ***_1_ − ***µ***_2_. These represent the separation between the homozygote clusters; if these are small, the homozygote clusters are not well separated.

2. ‖ ***µ***_1_ − ***µ***_2_. ‖_2_ If this is small, the homozygote clusters are not well separated.

3. The number of lines in the smaller of the homozygote clusters.

4. The number of lines in the error component.

Given multiple acceptable parameter estimates from multiple runs of the Markov chain, we select the estimates with the smallest value of max {|**∑**_1_|, |**∑**_2_|}. This heuristic is effective, because chains that have become stuck in a local maximum and not converged, generally (incorrectly) group multiple clusters into a single (excessively large) cluster. This results in covariance matrices with large determinants. Markers with more than two homozygous alleles must be analysed using some other method, e.g. DBSCAN [Ester et al., 1996].

## 4 Example

We applied this genetoype calling method to data for 90,000 markers in a wheat population of 3,412 lines, generated by an eight-parent MAGIC design [Shah et al., 2019]. We used 200 MCMC samples for each Markov chain, and ran eight Markov chains per marker. Our choice of hyperparameters was

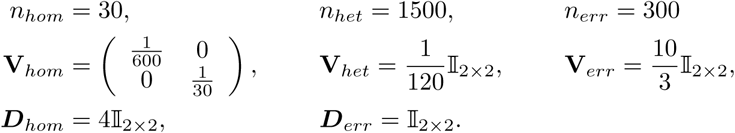

These parameters were chosen by checking their performance on a subset of 200 markers. These parameters capture prior information in the following ways:

1. Our choice of **V**_*hom*_ implies that homozygote clusters have more variability along the *r*-axis than the *θ*-axis, however the small value of *n*_*hom*_ implies that this information is weak.
2. Our choice of **V**_*het*_ implies that the heterozygote cluster is small. The large value of *n*_*het*_ implies a very strong prier belief.
3. The error cluster is large (flat distribution), with a moderate degree of certainty.

To achieve good results with few MCMC steps, the Gibbs sampler must have a sensible starting point. We initialized the *r* coordinates of ***µ***_1_ and ***µ***_2_ to 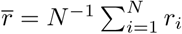. We considered a model acceptable if all the following held:

1. 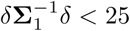 and 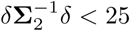, where *δ* = ***µ***_1_ − ***µ***_2_.
2. ‖***µ***_1_ − ***µ***_2_‖_2_ > 0.06.
3. Each homozygote cluster contained more than 200 lines.
4. The proportion of lines in the error component was smaller than 0.25.

The ‘best’ chain was selected as the chain with the smallest value of max {|**∑**_1_|, |**∑**_2_|}. A correctly called marker is shown in Figure 1. Of 90, 000 markers, only 6,743 needed to be manually reviewed. Some of these reviewed markers were determined to have more than two marker alleles, and were called using DBSCAN. Others were cases of MCMC non-convergence, for which the MCMC algorithm was run again.

